# RNAseq-based gene expression analysis of *Melolontha hippocastani* hindgut pockets and the surrounding hindgut wall tissue

**DOI:** 10.1101/2022.12.01.518689

**Authors:** Pol Alonso-Pernas, Wilhelm Boland

**Affiliations:** Department of Bioorganic Chemistry, Max Planck Institute for Chemical Ecology, Jena, Germany

## Abstract

In this study, the metatranscriptome of newly-discovered structures attached at the distal end of the hindgut of the larvae of a coleopteran (*Melolontha hippocastani*), is compared with that of the surrounding hindgut wall. Larvae were collected in their natural habitat, RNA was extracted using a commercial kit and sequenced in a Illumina HiSeq2500 platform. 250 bp paired-end reads were used to de novo assemble the transcriptomes. Contig annotation was carried out with BLASTx and Blast2GO PRO and differential expression analysis was performed in edgeR. Contigs aligned mainly to *Achromobacter* sp. in the pockets and to the Firmicutes phylum in hindgut wall. Host RNAs were expressed in the pockets in higher amounts than in hindgut wall. Gene expression suggest that pocket bacteria undergo aerobic metabolism and are exposed to higher levels of oxidative stress than the population of the hindgut wall. Hypothetical functions for the pocket might be immune-stimulation and regulation of host development, while the hindgut wall appears to be devoted to degradation of dietary polysaccharides and host nitrogenous wastes. Further research is necessary to experimentally prove these suggested roles.

## 1. Introduction

Insects are one of the most successful animal groups on earth and they thrive on a plethora of different diets. Although their digestive tracts present a wide variety of morphologies as result of adaptation to disparate food sources (Engel & Moran 2013) they are generally divided in the same three sections: the foregut, where the food is preprocessed and digestion may start; the midgut, where host digestive enzymes are secreted into the gut lumen and, in combination with symbiotic activity, most of the digestion and nutrient absorption takes place; and the hindgut, were water and other small molecules are absorbed (Chapman 2013). However, in some insect groups, this scheme can be subjected to significant modifications. For example, in termites or scarabaeid beetles, the hindgut is enlarged in the so called paunch (in termites) or fermentation chamber (in scarabaeids) and it also participates in digestion with the aid of symbiotic microorganisms (Calderon-Cortes et al. 2012). Or in *Spodoptera* larvae, the foregut is enlarged and accumulates α- and β-carotene, apparently as an adaptation to feeding on toxic plants (Shao et al. 2011).

In some cases, the morphological diversification of the digestive tract creates additional structures exclusive of particular insect taxa. These structures may be devoted to the housing of a specific bacterial ectosymbiont: for example, the midgut crypts in bugs of the Alydidae family are colonized by the environmental bacterium *Burkholderia* sp. (Kikuchi 2005) or by Gammaproteobacteria in bugs of the Pentatomidae and Cydnidae families (Prado & Almeida 2009; Hosokawa et al. 2012); also, the pouch-like cavity located in the midgut-hindgut junction of *Tetraponera* ants is occupied by root-nodule nitrogen fixing related bacteria (van Borm et al. 2002). In other insects, bacterial or fungal endosymbionts inhabit specialized cells (bacteriocytes or mycetocytes) which usually are grouped into clusters called bacteriome or mycetome. This is the case of the lygaeid *Kleidocerys resedae* or the chrysomelid *Bromius obscurus* that house gammaproteobacterial symbionts within, respectively, midgut or foregut-midgut junction associated bacteriomes (Küchler et al. 2010; Fukumori et al. 2017). Similarly, the cerambycid beetles *Tetropium castaneum*, *Rhagium inquisitor* and *Leptura rubra* harbour, respectively, unidentified yeast, *Candida rhagii* and *Candida shehatae* endosymbionts within mycetomes associated with the proximal midgut (Grünwald et al. 2009). The purpose of the microbes inhabiting such symbiont-containing organs is unclear in most of the cases, although nitrogen fixing has been suggested for the root nodule-related bacteria within the paunch of *Tetraponera* ants (Stoll et al. 2007) and *Burkholderia* sp. associated with the bean bug *Riptortus pedestris* has been proven capable of degrading the insecticide fenitrothion (Kikuchi et al. 2012) and promoting insect growth and egg production through modulation of host protein expression (Lee et al. 2017). Regarding the endosymbiotic microorganisms confined within bacteriomes or mycetomes, it is commonly accepted that they provide a variety of essential nutrients to the host (Bright & Bulgheresi 2010).

RNA sequencing (RNAseq) has been successfully applied to explore the gut function of insects under changing conditions (Roy et al. 2016) as well as the interaction of insect host with either symbiotic (Peterson & Scharf 2016; Emery et al. 2017) or pathogenic organisms (Chen et al. 2017; Yadav et al. 2017). Tissue-specific RNAseq can also unmask differentially expressed genes and functions that could otherwise remain unidentified in whole-gut analysis (Mamidala et al. 2012; Shelomi 2017; Nakayama et al. 2017). In the present experiment, we used Illumina Hiseq to survey the transcriptome of the hindgut pockets of the forest cockchafer (*Melolontha hippocastani*). These symbiont-containing organs, placed at both sides of the terminal segment of the larval (male and female) hindgut chamber, were discovered in the 50s (Wildbolz 1954) and their architecture and bacterial community have been recently characterized in detail utilizing modern techniques (Alonso-Pernas et al. 2017). However, little is known about the processes ongoing in these enigmatic structures, such as the mechanism of symbiotic colonization or their role in the context of insect physiology. We expect to get an insight into these matters by comparing the pocket transcriptome with that of the surrounding hindgut wall of larvae retrieved straight from their natural environment.

## 2. Materials and methods

### 2.1. Insect collection and RNA extraction

Second-instar (L2) *Melolontha hippocastani* larvae were collected from a deciduous forest next to Pfungstadt (Germany, 49°49’44” N 8°36’17” E) in June 2016. Larvae were carried to the laboratory in their natural soil and flash-frozen with liquid nitrogen immediately after arrival in order to minimize changes in gene expression. Frozen larvae were stored at −80 °C until further processing. Diethyl pyrocarbonate (DEPC, Sigma Aldrich, Saint Louis, MO, USA) treated phosphate buffered saline solution (PBS) and ddH_2_O (Nagy et al. 2007) were prepared by adding 0.1% v/v DEPC into PBS (composition per liter: 8 g NaCl, 0.2 g KCl, 1.44 g Na_2_HPO_4_, 0.24 g KH_2_PO_4_ (pH 7.4)) or ddH_2_O and incubating the mixtures at 37 °C for 24h before autoclaving. Dissection of thawed insects was carried out on ice in DEPC-treated PBS using ethanol sterilized tools rinsed with DEPC treated ddH_2_O. Entire guts were carefully pulled out from the larvae. Pockets and 1×1 mm pieces of nearby hindgut wall were excised from the hindgut chamber and rinsed in DEPC-ddH_2_O to remove debris and unattached bacteria. Tissues were pooled together in a 2 mL polypropylene screw cap micro tube (Sarstedt, Nümbrecht, Germany) forming 3 hindgut wall replicates, each consisting of 8 hindgut wall fragments from 8 different individuals, and 1 pocket replicate, consisting of approximately 100 pockets of 50 different individuals. Reduced size of pocket tissue prevented the obtainment of more replicates. Tubes containing dissected tissues were kept in liquid nitrogen until pooling was completed. RNA extraction was performed using the innuPREP RNA Mini Kit (Analytik Jena, Jena, Germany) with an extra homogenization step at the beginning of the protocol: 450 μl of Lysis Solution RL, 6 μL of lysozyme solution (50 mg/mL) and bashing beads were added to the sample 2 mL PP micro tube before homogenization with a 2010 Geno/Grinder^®^ device (Spex^®^SamplePrep, Metuchen, NJ, USA) during 1 min at 1250 rpm. Samples were incubated for 3 min at RT and then homogenized again under abovementioned conditions. Rest of RNA extraction was conducted according to manufacturer’s instructions. RNA quality and purity was tested with a Experion^™^ RNA chip (Bio-Rad, Hercules, CA, USA) on a 2100 Bioanalyzer (Agilent, Santa Clara, USA) following manufacturer’s protocols.

### 2.2. RNA sequencing and de novo transcriptome assembly

RNA samples were sent on dry ice to the Max Planck Genome Centre in Cologne (Germany) for sequencing on an Illumina HiSeq2500 platform. Prior to library preparation ribosomal RNA was depleted from the samples. Random primed cDNA libraries were prepared with a TruSeq RNA Sample Prep kit (Illumina, San Diego, CA, USA) and sequenced, yielding 8 million (4 gigabases, each hindgut wall library), and 12 million (6 gigabases, pocket library) of 250 bp paired-end reads. Library quality control, adaptor trimming and *de novo* transcriptome assembly were carried out using CLC Genomics Workbench (Qiagen, Redwood City, CA, USA). A reference transcriptome was generated by a combined assembly of both hindgut wall and pocket reads using CLC Genomics Workbench v10.1 software with standard settings and an additional CLC-based assembly with the following parameters: word size = 64, bubble size = 300; minimum contig length = 350 bp; nucleotide mismatch cost = 2; insertion = deletion costs = 2; length fraction = 0.7; similarity = 0.9. Conflicts among individual bases were resolved in all assemblies resolved by voting for the base with the highest frequency. Contigs shorter than 350 bp were removed from the final analysis. The two assemblies were compared according to quality criteria such as N50 contig size, total number of contigs and the number of sequence reads not included in the contig assembly. For each assembly, the 50 largest contigs were manually inspected for chimeric sequences. The presumed optimal consensus transcriptome was selected from the different assemblies based on the highest N50 contig size, lowest total number of contigs and lowest number of chimeric sequences in the 50 largest contigs. The parameters of contig assembly were the following: word size = automatic, bubble size = 300; minimum contig length = 350 bp; nucleotide mismatch cost = 2; insertion = deletion costs = 2; length fraction = 0.7; similarity = 0.9. The resulting final *de novo* reference assembly (backbone) contained 305,905 contigs (minimum contig size = 350 bp) with an N50 contig size of 1,037 bp and a maximum contig length of 32,815 bp. The raw sequencing data was deposited in the NCBI Sequence Read Archive under the accession numbers SRR6123467 (hindgut wall replicate 1 sequencing run 1), SRR6123468 (hindgut wall replicate 1 sequencing run 2), SRR6123465 (hindgut wall replicate 2), SRR6123466 (hindgut wall replicate 3) and SRR6123464 (pockets).

### 2.3. Annotation and differential expression analysis

BLAST searches were conducted on a local computer cluster against the NCBI nr database with BLASTx (e-value cut-off 1e-1) and saved as .xml files. Further transcriptome annotation was carried out with Blast2GO PRO (Conesa & Stefan 2008) using Gene Ontology terms and EC numbers. Digital gene expression analysis was carried out using CLC Genomics workbench v10.1 to generate BAM mapping files, and finally by counting the sequences to estimate expression levels, using previously described parameters for read mapping and normalization. For read mapping, we used the following parameters: read assignment quality options required at least 60 % of the bases matching (the amount of mappable sequence as a criterion for inclusion) the reference with at least 94% identity (minimum similarity fraction, defining the degree of preciseness required) within each read to be assigned to a specific contig; mismatch cost = 2; insertion = deletion cost = 3; maximum number of hits for a read (reads matching a greater number of distinct places than this number are excluded) = 10; n-mer repeat settings were automatically determined and other settings were not changed. RPKMs (reads per kilo base per million mapped reads) (Mortazavi et al. 2008) were calculated from the raw read count to normalize the expression of each contig. Contigs with RPKM < 1 in all libraries were considered not transcriptionally active and were discarded from further analysis. Differential gene expression analysis between pocket and hindgut wall libraries was carried out using the R package edgeR (Robinson et al. 2010), using TMM (trimmed mean of M-values) normalized values of read CPM (counts per million) per contig (Oshlack & Wakefield 2009). Significance threshold of FDR (false discovery rate) corrected p value was set to 0.05. Since lack of replicates in one of the samples analyzed (pocket library) reduces statistical reliability in differential expression testing, the following components were included in the analysis: a) before TMM normalization, contigs with =< 1 CPM (counts per million) in all libraries were filtered out, regardless of their RPKM value. Therefore, genes with very low counts that may influence the accuracy of multiple testing were removed while maintaining tissue-specific expression patterns. b) Besides a FDR value =< 0.05, the requirement of fold change (FC) value > 10 or < −10 was imposed to define differentially expressed genes, thus discarding genes with low FDR value due to consistency among hindgut wall libraries but with little difference between hindgut wall and pockets. Additionally, in order to discriminate the major processes ongoing in pocket and hindgut wall, highly and differentially expressed (HDE) genes were selected by defining high expression as any RPKM 10 times above the library mean. Hindgut wall contigs were considered highly expressed when meeting such requirement in all three hindgut wall libraries. The web-based KEGG pathway mapping tool Search Pathway (http://www.genome.jp/kegg/tool/map_pathway1.html) was used to functionally annotate differentially expressed (DE) enzymes with associated EC code and to reconstruct metabolic pathways according to the Kyoto Encyclopedia of Genes and Genomes (KEGG).

## 3. Results

### 3.1. de novo transcriptome assembly and annotation

A total of four sequencing libraries (three for hindgut wall, one for pocket) were produced from larval *M. hippocastani* RNA. Each hindgut wall library contained, approximately, 8 million reads (4 gigabases). Each pocket library contained about 12 million reads (6 gigabases). Reads were assembled into a total of 305.905 contigs (Tables 1 and 2). After read alignment and RPKM normalization, contigs with very low expression (RPKM < 1 in all libraries) were discarded. The taxonomic assignment of contigs performed via Blast2GO PRO revealed that the most common eukaryotic alignment was the insect *Tribolium castaneum* (fam. Tenebrionidae) in both hindgut wall and pocket libraries, while the most common prokaryotic alignments were *Achromobacter* sp. (fam. Alcaligenaceae) in the pocket library and *Clostridium* sp. (fam. Clostridiaceae) in hindgut wall libraries (Fig. 1).

**Figure 1.**
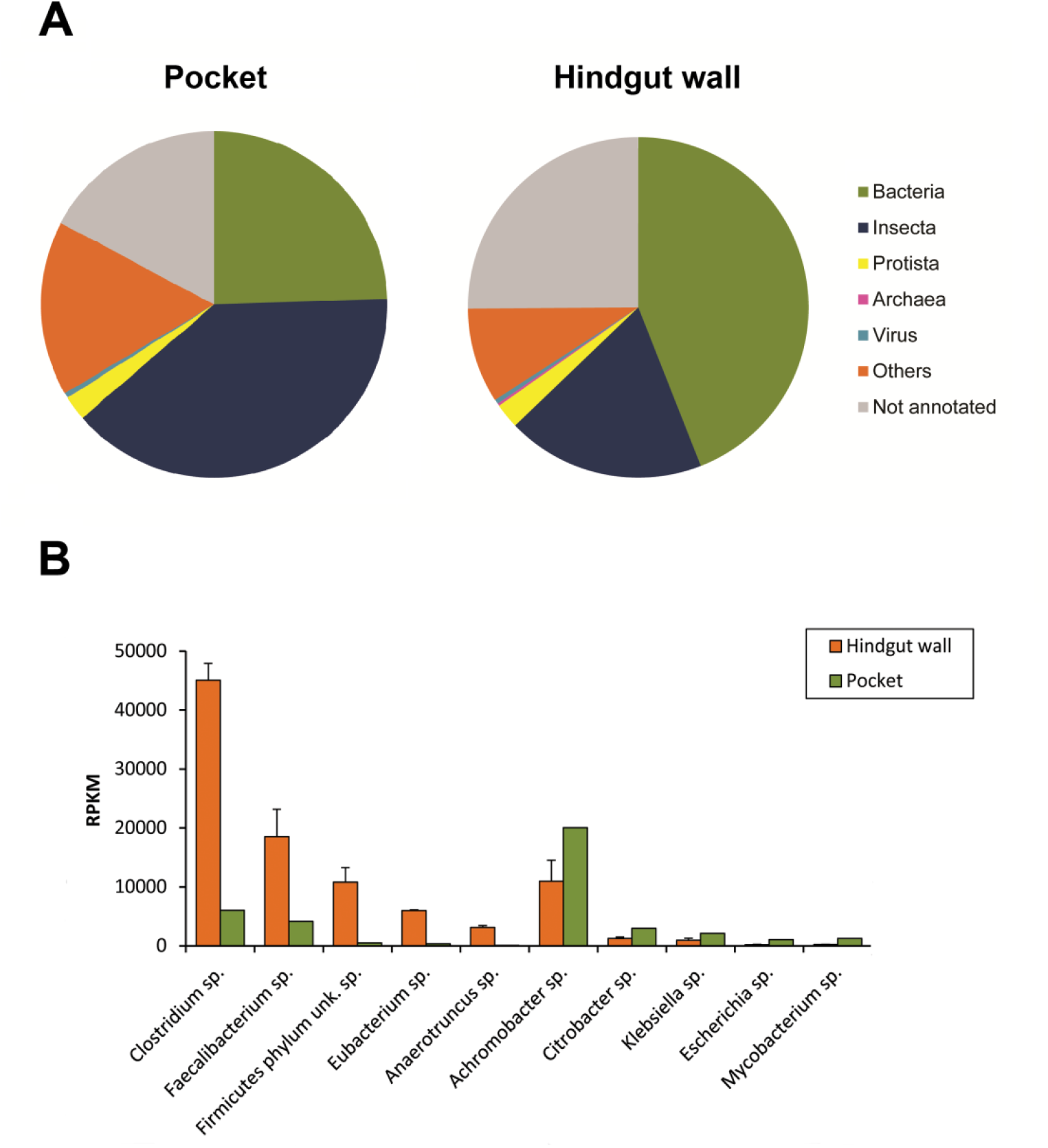
Taxonomic distribution of annotated contigs. After RPKM normalization, contigs with RPKM < 1 in all libraries were discarded. A) Pie charts showing the distribution of contigs in taxonomic groups. Hindgut wall chart represents the mean of the 3 hindgut wall libraries. B) Bar chart showing the five bacterial genera with higher RPKM in hindgut wall libraries (left half) and pocket library (right half). Orange bars represent the mean of all three hindgut wall libraries. Error bars represent standard deviation.

**Table 1.**
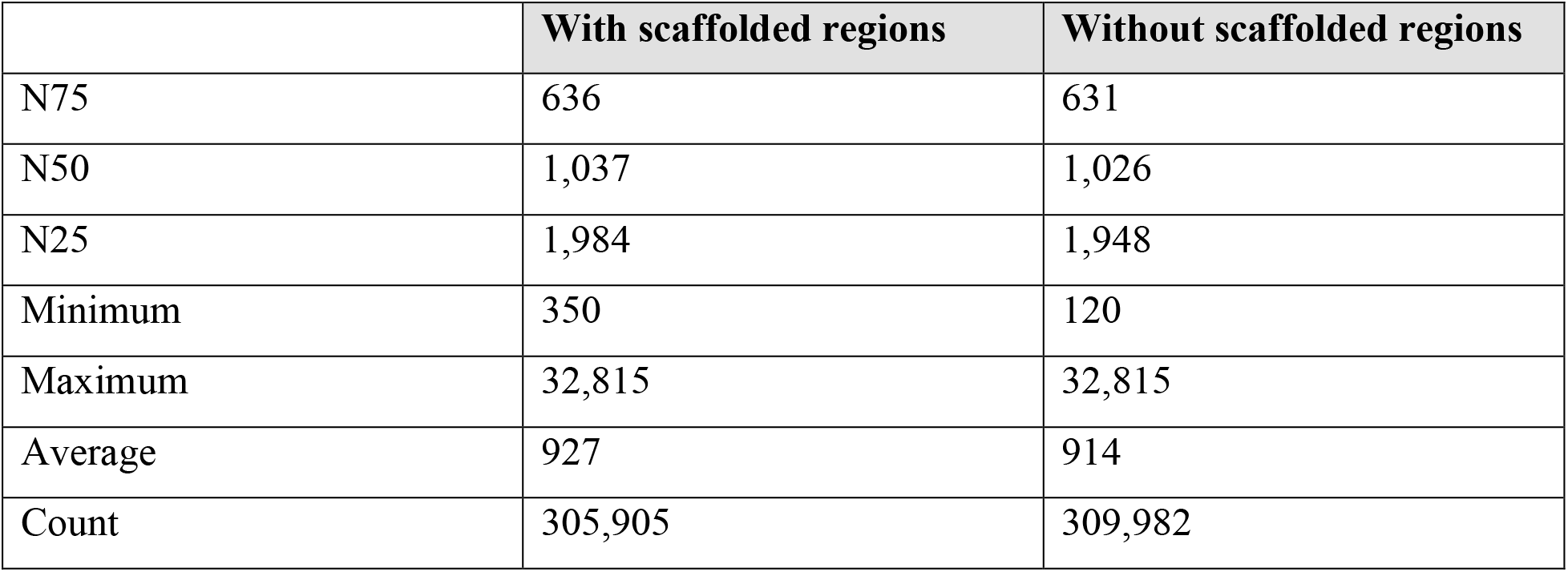
Contig length measurements

**Table 2.** Transcriptome assembly statistics

|  | Count | Average length | Total bases |
| --- | --- | --- | --- |
| Reads | 64,067,071 | 227.73 | 14,590,016,540 |
| Matched | 51,171,136 | 227.1 | 11,620,962,785 |
| Not matched | 12,895,935 | 230.23 | 2,969,053,755 |
| Reads in pairs | 43,000,840 | 273.87 | 283,424,930 |
| Broken paired reads | 8,102,050 | 244.72 |  |
| Contigs | 305,905 | 926 |  |

### 3.2. Differential expression analysis

After filtering contigs with CPM =< 1 in all libraries, a total of 105,693 contigs were considered for differential expression analysis. Of these, edgeR identified 23,856 differentially expressed (DE) contigs (FDR<0.05, FC > 10 or < −10) between hindgut wall and pocket libraries. Differentially expressed KEGG pathways in pocket and hindgut wall were reconstructed by the web-based tool Search Pathway using as input contig-associated EC codes. A summary of the top DE KEGG pathways is shown in Fig. 2. A comprehensive list of all DE contigs and the KEGG pathways to which they map is shown in Supplementary Table 1. Additionally, any contig with RPKM ten times above the mean of the library was considered as highly expressed. By applying this rationale, 1,106 contigs were found to be highly expressed in all three hindgut libraries, while 2,657 contigs were found to be highly expressed in the pocket library. By combining these statistics, 495 contigs were found to be highly and differentially expressed (HDE) in the hindgut wall compared to pocket, and 220 were found to be HDE in the pocket compared to hindgut wall (Fig. 3). The top identified HDE contigs in pocket and hindgut libraries are shown in Table 3. A comprehensive list of all HDE contigs and the KEGG pathways to which they map is shown in Table S2, and can be summarized as follows: unique HDE genes in pocket library were, among others, 20 contigs that mapped to insect cuticular proteins and precursors, 15 to eukaryotic transposases and transposable elements, 9 to insect chitinases and chitin-binding proteins, 8 to bacterial porins, 6 to eukaryotic reverse transcriptases, 6 to insect CD109 antigen, 4 to putative eukaryotic growth factors (3 of which are described as multiple epidermal growth factor-like domains), 4 to putative insect sulfate transmembrane transporters (sodium-independent sulfate anion transporter-like), 4 to eukaryotic histone-lysine N-methyltransferase and 3 histone-lysine N-methyltransferase-like, 3 to insect glucose dehydrogenases, 3 to insect Osiris proteins and precursors, 2 to toxic bacterial proteins of Hok family, 2 to insect yellow protein, 2 to proteins involved in insect response to stimulus (cuticular analogous to peritrophins 1-J isoform X2 and odorant binding 4), 2 to insect peroxidases and one to a bacterial peroxiredoxin, an insect serine protease H164 (EC:3.4.21), a bacterial lipoyl synthase (EC:2.8.1.8), an insect phenoloxidase subunit A3, a bacterial genetic competence-related protein, an insect arylphorin subunit alpha, a bacterial Tu translation elongation factor, an insect C-type lectin and an insect gram-negative bacteria binding protein. Unique HDE genes in hindgut libraries were, among others, 10 contigs that mapped to bacterial flagellar proteins, 10 to bacterial carbohydrate ABC transporters and transcriptional regulators, 8 to bacterial proteases (6 of which are serine-type (EC:3.4.21; EC:3.4.17; EC:3.4.16)) and regulators, 7 to bacterial sporulation-related proteins, 6 to bacterial chemotaxis-related proteins, 5 to bacterial amino acid transmembrane transporters, 5 to bacterial proteins of pF06949 family, 4 to bacterial collagen repeats, 3 to bacterial aldehyde oxidoreductases (EC:1.2.99.7), 3 to bacterial aldehyde ferredoxin oxidoreductases (EC:1.2.7.5), 3 to insect cystathionine beta-synthases, 2 to bacterial carbamate kinases (EC:2.7.2.2), 2 to bacterial C4-dicarboxylate ABC transporters, 2 to bacterial iron transmembrane transporters, 2 to bacterial alkaline-shock proteins, 2 to bacterial beta lactamases, 2 to bacterial pyrophosphatases, 2 to bacterial addiction module toxin system, one to an insect aquaporin AQPcic, a bacterial ferredoxin, a bacterial manganese-containing catalase, a negative regulator of bacterial genetic competence, a bacterial citrate transporter and an insect histone deacetylase complex subunit SAP18.

**Figure 2.**
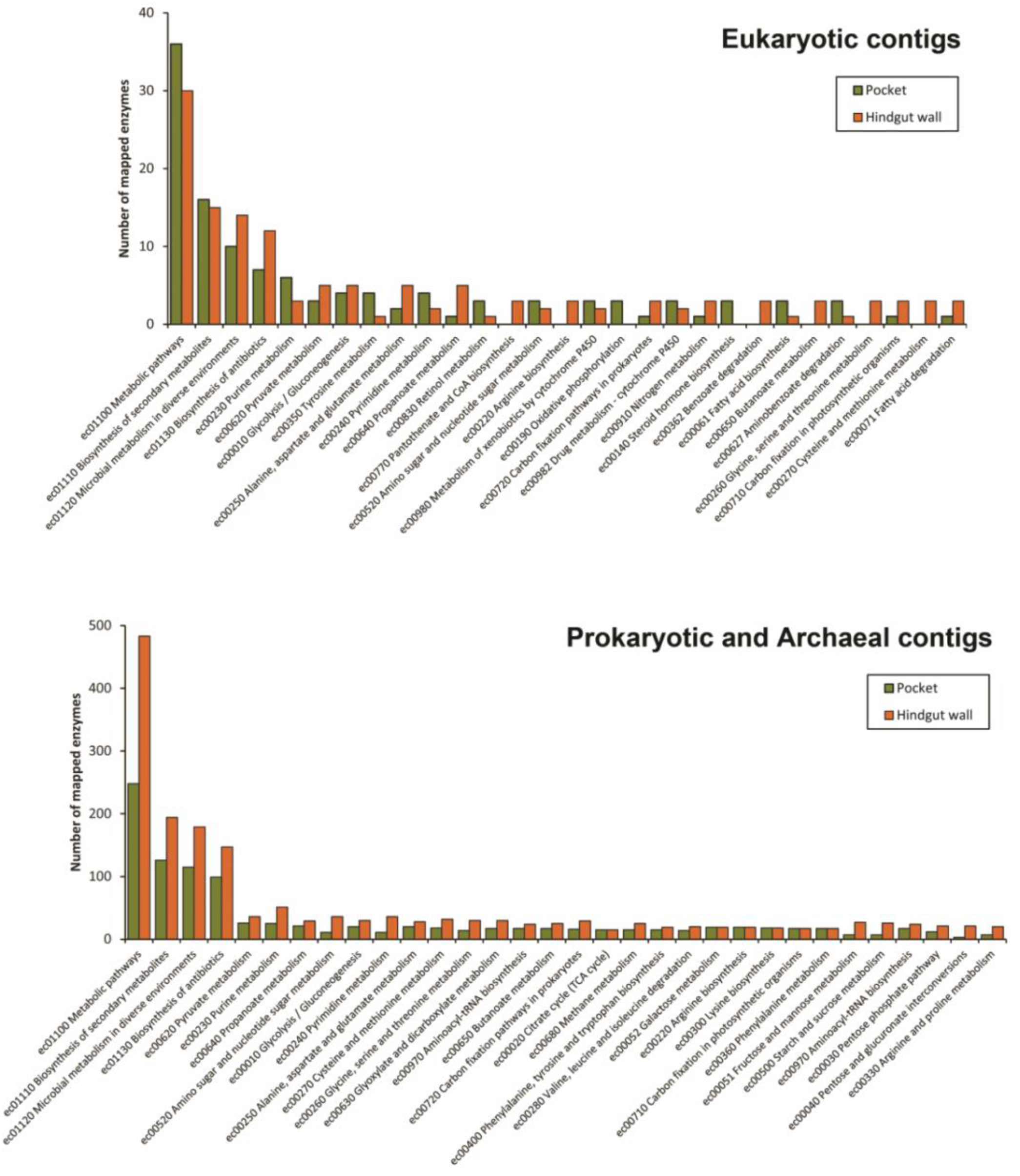
Top DE KEGG pathways in pocket and hindgut wall tissues sorted according to contig taxonomic assignment.

**Figure 3.**
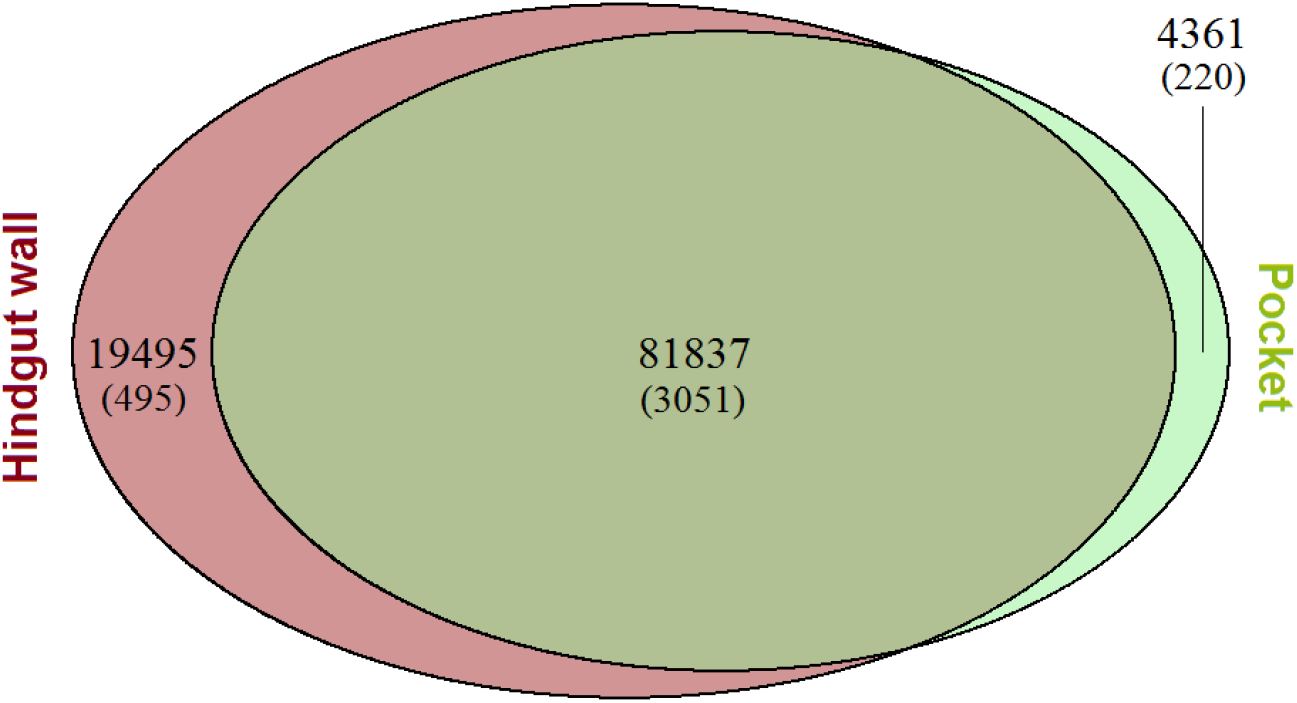
Venn diagram showing the distribution of contig expression between libraries. Brown area shows DE sequences in hindgut wall, green area shows DE sequences in pocket. Overlapping area shows sequences equally expressed in both tissues. Number of sequences (in parentheses, highly expressed sequences) contained in each area are given.

**Table 3.**
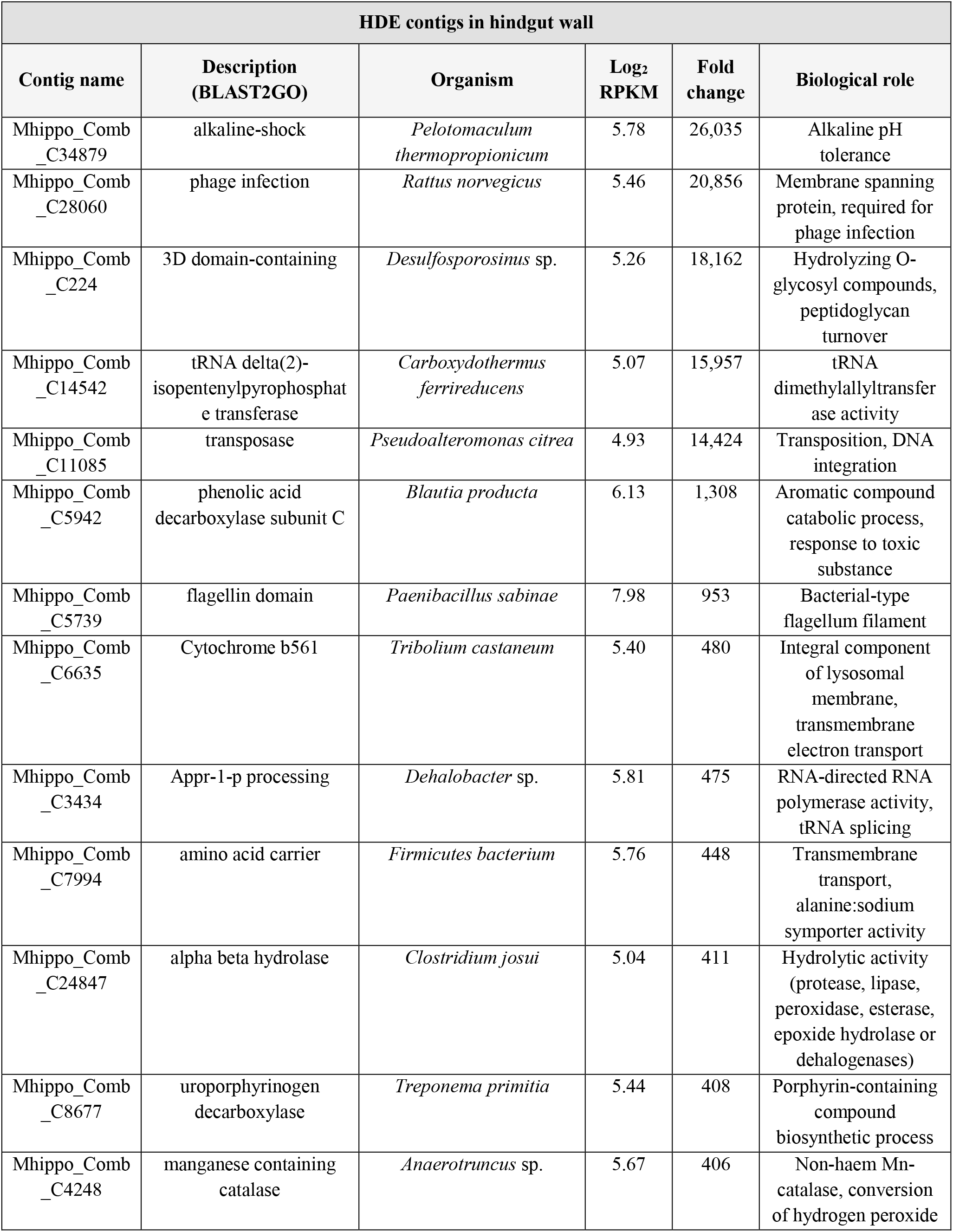
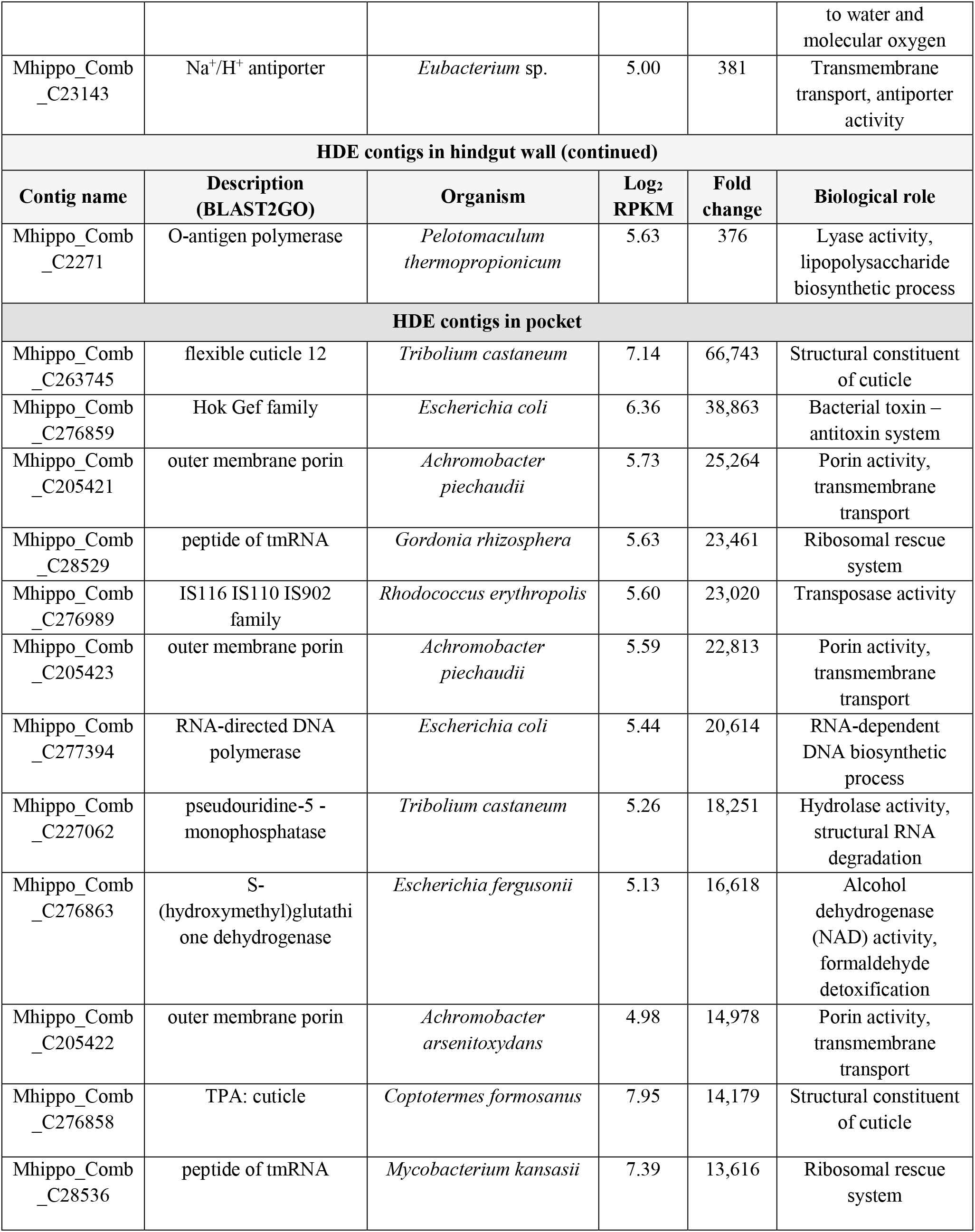
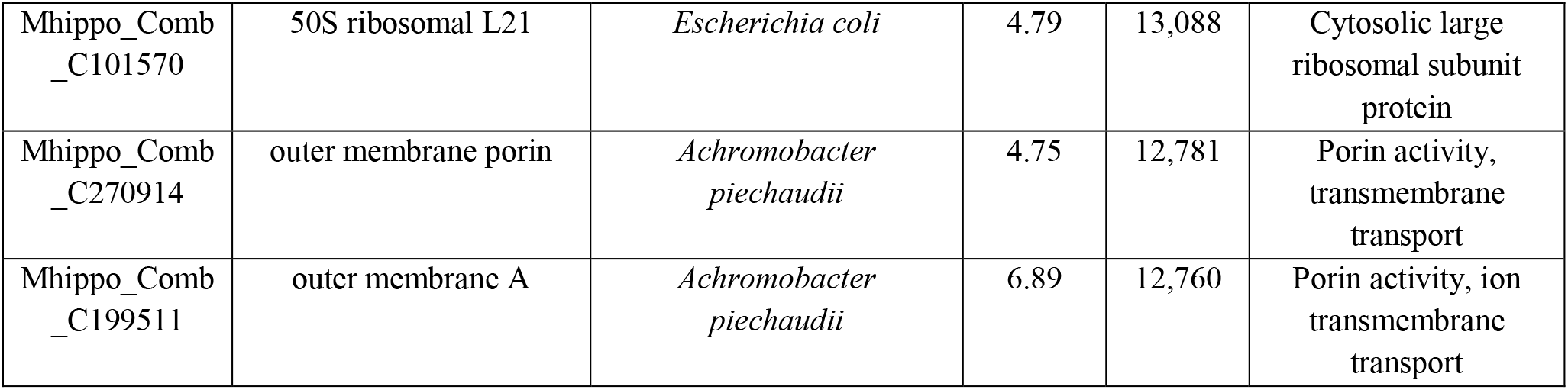
Top identified HDE contigs in hindgut wall and pocket, sorted in decreasing order of fold change. Only contigs highly (RPKM > 10× library mean) and differentially expressed (FDR < 0.05, FC > 10 or < −10), annotated to proteins with known function are shown. In case of hindgut wall, log_2_ RPKM column shows mean of log_2_ transformed RPKMs of the three libraries. Fold change column shows the RPKM fold change compared to the other tissue.

## 4. Discussion

### 4.1. Taxonomic distribution of reads

As shown in Fig. 1A, the majority of annotated reads in pocket and hindgut wall fell into insect or bacterial taxa. Protist-annotated reads were also detectable in both tissues, although they are likely to originate from transient or parasitic microorganisms or from wrongly annotated insect-derived RNAs, as their symbiotic association with terrestrial coleopterans is exceptional (Tanahashi et al. 2017). Archaeal reads were also detected in both tissues, although in negligible amounts (less than 0.5%) which is in concordance with previous observations in *Melolontha melolontha* (Egert et al. 2005). Finally, around 10 and 20% of reads in hindgut wall and pocket tissue, respectively, mapped to non-insect eukaryotic organisms. Considering the lack of reference genome for *Melolontha hippocastani* and the inherent alignment uncertainty, all eukaryotic-mapped reads were assumed to be of host origin. Bacteria-mapped contigs in pocket and hindgut wall libraries fell into different genera depending on the tissue (Fig. 1B). In hindgut wall, reads mapped mostly to anaerobic gram-positive bacteria, while in the pocket they mapped mostly to aerobic or facultative anaerobic gram-negative bacteria. Differences in community composition between pockets and hindgut wall were previously observed; however, the Clostridiales and the Clostridiaceae, the order and family with highest number of mapped reads in the hindgut wall, were only poorly detected in the hindgut wall of second-instar larvae (Alonso-Pernas et al. 2017). They were, nevertheless, very abundant in third-instar larvae. This suggests that, although the larvae used in the present study had the body size corresponding to second-instar larvae, the composition of their gut microbiome was already drifting to third-instar’s. Following this hypothesis, negligible number of reads mapped to the Micrococcaceae family in pocket libraries, although they were very abundant in second-instar larvae (Alonso-Pernas et al. 2017). This suggests that this bacterial family is either dormant or ultimately outgrown by the Alcaligenaceae family, concretely by the genus *Achromobacter*.

### 4.2. Highly and differentially expressed (HDE) genes in hindgut wall

The disparate annotation of highly and differentially expressed (HDE) contigs in pockets and hindgut wall is indicative of functional differences between the two tissues. In hindgut wall, a bacterial alkaline shock gene showed the highest fold change compared to pockets, which is in line with the alkaline pH (around 8) of this section of the gut (Egert et al. 2005) (Table 3). Also, many bacterial genes related to sporulation were HDE in hindgut wall (Table S2), which is not surprising due to the high abundance of Clostridial RNAs in the libraries. Sporulation-related genes are commonly expressed in gram-positive symbionts of insects and other organisms (Margulis et al. 1998; Paredes-Sabja et al. 2011) as a response to biofilm formation and high bacterial density (Dapa et al. 2013; McKenney et al. 2013). Other HDE genes suggesting ongoing bacterial colonization of the hindgut wall and microbe-microbe interactions were flagellar and chemotaxis-related contigs (Rawls et al. 2007; Wang & Wood 2011; Stephens et al. 2015), the toxin-antioxin system addiction module (Wang & Wood 2011), bacterial collagen repeats that might anchor bacteria in the gut wall and participate in host-microbe interactions (Yu et al. 2014) and betalactamases that may protect bacteria from antibiotics secreted by neighbouring microorganisms (Chen et al. 2017) or from toxic dietary compounds (Allen et al. 2009). All these HDE genes related to symbiotic colonization and interactions indicate that the hindgut wall community of *M. hippocastani* is not stable but subjected to continuous changes, which agrees with previous observations (Alonso-Pernas et al. 2017). HDE bacterial aldehyde oxidoreductases might neutralize antimicrobial agents contained in the diet (Correia et al. 2016). HDE bacterial transcripts for iron transporters were possibily a consequence of limitation of this element, whose availability in the gut is tightly controlled by the insect (Nichol et al. 2002). Congruent with iron scarcity is the presence of a HDE bacterial manganese-containing catalase transcript, a non-heme catalase expressed under conditions of microaerophilic oxidative stress and iron depletion (Whittaker 2012). Many ABC transporters and transcriptional regulators involved in transmembrane transport of carbohydrates were also HDE by hindgut wall bacteria, reflecting their role, along with insect secreted enzymes, in the digestion of recalcitrant polysaccharides, which are degraded extracellularly and the resulting soluble saccharides are internalized for further processing (Martin 1983; Artzi et al. 2017). HDE bacterial protease transcripts (mostly serine proteases) and the alkaline pH occurring in the hindgut, which might contribute to the solubilization of dietary proteins, supports the digestive role of the resident community (Vinokurov et al. 2006). HDE bacterial amino acid transmembrane transporters (most of which were identified as sodium:solute symporters) suggest occurrence of free amino acids, probably due to ongoing proteolysis or *de novo* synthesis from host nitrogenous waste (Alonso-Pernas et al. 2017). Differentially expressed (DE) urease (EC:3.5.1.5) and carbamate kinase (EC:2.7.2.2.) in hindgut wall support presence of ammonia in hindgut (Table S1), as the former produces it by hydrolyzing urea and the latter uses it as substrate. HDE insect cystathionine beta-synthases (EC:4.2.1.22), which plays a role in the synthesis of the essential amino acid cysteine, might be related to the production of cysteine-rich proteins that regulate symbiotic population (Futahashi et al. 2013). HDE insect aquaporin might be involved in water absorption from feces, a well-reported hindgut function (Campbell et al. 2008; Chapman 2013). Finally, HDE transcripts for insect histone deacetylase complex may suggest repression of host genes, which is in line with the low number of detected host RNAs (Fig. 1) (Martin & Zhang 2005; Haberland et al. 2009). Alternatively, insufficient sequencing depth might have caused masking of low-abundant insect RNAs by highly-abundant bacterial RNAs, in which case some of the host’s functions may have gone unnoticed.

### 4.3. Highly and differentially expressed (HDE) genes in pocket

An obvious increase in host-related HDE transcripts was observed in the pocket library compared to hindgut wall, possibly due to histone-lysine N-methyltransferase mediated upregulation (Table S2) (Martin & Zhang 2005). Most of the host HDE transcripts in pocket were cuticle-related. Insect cuticle constitutes a defensive barrier in the external body as well as in foregut and hindgut (Chapman 2013). Among pocket HDE transcripts both insect structural constituents of cuticle and chitinases are found, which suggest simultaneous cuticle synthesis and degradation. This may indicate either cuticular renovation, as the old cuticle must be degraded before being replaced by the new one (Willis et al. 2012; Chapman 2013), or cuticle thickening as a response to external challenge (Li & Denlinger 2009; Bascuñán-García et al. 2010). The latter possibility is supported by early observations of the tissue (Wildbolz 1954) and by the presence of HDE eukaryotic epidermal growth factors. It has been shown that other functions of these proteins are protecting the integrity of the intestinal barrier, reducing colonization by pathogens and attenuating the epithelial inflammatory response (Tang et al. 2016; Kim et al. 2016). Further hints pointing towards active host immune response in pocket tissue are the HDE insect transcripts for gram-negative bacteria binding protein, C-type lectin and HDE CD109 antigen-like proteins which might be involved in pathogen recognition and activation of host defenses (Kim et al. 2000; Zelensky & Gready 2005; Warr et al. 2008; Yazzie et al. 2015), for a phenoloxidase domain and a serine protease, both possibly involved in melanin synthesis (Ashida & Breyt 1995) and for a yellow protein, which positions the melanin pigment in the cuticle (Willis et al. 2012) among other defensive roles (Gretscher et al. 2016). Melanin synthesis in pockets is further supported by the fact that in some individuals the pockets appear stained black (Wildbolz 1954). Functions of this pigment are mostly defensive, i.e. wound healing or pathogen encapsulation (Nakhleh et al. 2017). This points towards a tight regulation of the pocket population by the innate immunity of the host which, in combination with the oxidative stress suggested by HDE insect peroxidases and bacterial peroxiredoxins (Lushchak 2010), may have the purpose of preventing hindgut anaerobic bacteria from colonizing the pockets. Additionally, host proteases may directly control symbiotic titer (Byeon et al. 2015) and yellow proteins appear to be up- and downregulated depending on seasonal conditions (Daniels et al. 2014; Vilcinskas & Vogel 2016) raising the question of whether seasonal changes have an influence in pocket gene expression. The high number of HDE contigs that mapped eukaryotic transposases, transposable elements and reverse transcriptases might be consequence of oxidative stress (Giorgi et al. 2011) or of host perception of pocket colonization as an infection event (Mhiri et al. 1997). In view of the high number of pocket reads assigned to *Achromobacter* sp. (Fig 1B), it is tempting to assume that host immune system is less detrimental to this genus than to the others. The mechanisms underlying this possible selectiveness are to be studied, but is it plausible that HDE insect C-type lectin is involved, since lectins can be used by certain bacteria to evade host anti-microbial peptides (Pang et al. 2016). Also, the presence of symbiotic *Achromobacter* in the pocket might stimulate host immune system as reported in other systems, thus conferring resistance against pathogens (Kim et al. 2016). HDE contigs mapping to porins were also very abundant in the pocket. These passive transporters are the most abundant proteins in the outer membrane of gram-negative bacteria and their high expression in pockets suggests an intense exchange of nutrients and ions between symbionts and environment (Galdiero et al. 2012). Interestingly, porin-assigned reads aligned mainly to *Achromobacter* sp., which supports a significant role for this bacterium in the pocket. Porins are crucial for the symbiotic colonization of the squid light-emitting organ (Aeckersberg et al. 2001), for the infection of insect hemocoel by an entomopathogen and possibly for the colonization of the gut of its nematode vector as well (Van Der Hoeven & Forst 2009). Moreover, porins have been related to invasion of non-insect tissues and manipulation of host defenses by pathogenic bacteria (Provenzano & Klose 2000; Duperthuy et al. 2011). Collectively, these observations suggest that *Achromobacter* sp. is engaged in a porin-mediated colonization process that may contribute to the triggering of insect immune response. Unfortunately, considering the current data any speculation on the role of the pockets as a whole is still far-fetched. Nevertheless, previous reports (Alonso-Pernas et al. 2017) and the presence of HDE sulfate transmembrane transporters suggests that the pockets might participate in nutrient provisioning. It is also noteworthy the pocket HDE contigs mapping to the Osiris gene family. These proteins are exclusive of insects, and although their role is unclear, they show expression peaks at specific life stages (embryo, second instar larvae and pupae) and might be involved in insect development (Shah et al. 2012). Also the HDE insect arylphorin subunit alpha and glucose dehydrogenases point towards the pockets being involved in host development. Alrylphorins are ubiquitous proteins in *Melolontha* sp. and other insects that act as amino acid storage proteins. Their concentration in hemolymph increases during each larval stage to quickly drop at each molt, as their amino acid components are used for the synthesis of body tissues (Delobel et al. 1993; Chapman 2013). Expression of glucose dehydrogenases is highly correlated with that of 20-hydroxyecdysone, a major insect molting hormone, and is increased during metamorphosis (Cox-Foster et al. 1990). HDE arylphorins, Osiris family proteins and glucose dehydrogenases make tempting to speculate that the role of the pockets and, perhaps, their bacterial symbionts, might go beyond insect immunity and nutrition and embrace other aspects of physiology such as development, as it happens in the *Riptortus-Burkholderia* system (Lee et al. 2017) or *Aedes* mosquitoes (Coon et al. 2017).

### 4.4. Differentially expressed (DE) metabolic pathways in hindgut wall

Taking into consideration all differentially expressed (DE) enzymes (FDR<0.05, FC>10 or <-10) with associated EC codes, DE KEGG pathways in pocket and hindgut libraries were reconstructed. The taxonomic assignment of the BLAST top hit allowed separation in host- and symbiont-derived pathways (Fig. 2, Table S1). Symbiotic DE enzymes belonging to the KEGG pathways purine and pyrimidine metabolism, and aminoacyl tRNA biosynthesis were DE in both pocket and hindgut wall libraries. This indicates ongoing DNA, RNA and protein synthesis, thus high symbiont growth and activity in both tissues. Within these pathways, allantoinases (EC 3.5.2.5), allophanate hydrolases (EC 3.5.1.54) and ureases (EC 3.5.1.5) were DE only in hindgut wall, suggesting degradation of insect nitrogenous waste, that is, uric acid and urea, to ammonia, as part of a nitrogen recycling mechanism (Alonso-Pernas et al. 2017). Proximity of the outlet of the Malpighian tubules, located in the midgut-hindgut junction, to the site of sampling of hindgut wall tissue supports this hypothesis, as Malpighian tubules collect hemolymph waste and pour it into the hindgut (Shelomi 2017). On the contrary, the glutamine synthetase pathway was DE by both pocket and hindgut wall symbionts, suggesting that pocket bacteria utilize hindgut-produced ammonia for the synthesis of amino acids. Symbiotic chitinases were DE only in hindgut wall, probably reflecting the usage of this cuticular polysaccharide by hindgut bacteria as nitrogen and carbon source, thus incidentally contributing in maintaining the optimal thickness of the intima layer for proper diffusion of nutrients (Indiragandhi et al. 2007). In the hindgut wall more DE symbiotic enzymes involved in KEGG pathways related to carbon metabolism compared to pockets were found (fructose and mannose metabolism and starch and sucrose metabolism). This is in line with the digestive role of hindgut bacteria discussed above. Cellulases were only DE by hindgut wall bacteria. Taken together, these observations indicate that pocket bacteria may not be directly involved in digestion of insect food. Symbiotic enzymes belonging to carbon fixation KEGG pathways such as the reductive TCA cycle and the Wood-Ljungdahl pathway were DE in hindgut wall. They may contribute to the supply of acetyl-CoA needed for bacterial metabolism, but also to the production of acetate which might be taken up by the insect. Bacterial acetogenesis is well documented within the digestive tract of wood-feeding termites and roaches (Warnecke et al. 2007; Zhang et al. 2010) as well as in other insects with disparate diets (Matson et al. 2011). High acetate concentration in the gut fluid of *Melolontha melolontha* and hydrogen accumulation in the midgut, but not in the hindgut, is in line with ongoing acetogenesis in the hindgut, as this process uses hydrogen as electron donor, thus preventing its accumulation (Egert et al. 2005).

Hindgut wall symbionts might also provide to the host, nutrients such as niacin, pantothenic acid, biotin and pyridoxine, as suggested, respectively, by the DE KEGG pathways nicotinate and nicotinamide metabolism, pantothenate and CoA biosynthesis, biotin metabolism and vitamin B6 metabolism (Gilmour 1961; Cohen 2015). Methanogenesis might happen in hindgut wall as well, as coenzyme-B sulfoethylthiotransferase (EC 2.8.4.1) is DE in libraries of this tissue. However, this is likely to be a minor process carried out by the small archaeal population (Fig. 1, Table S1) (Egert et al. 2005). Finally, the pathways yielding geranyl pyrophosphate, geranylgeranyl pyrophosphate and farnesyl pyrophosphate from pyruvate (within the KEGG pathway terpenoid backbone biosynthesis) were DE by hindgut wall symbionts, suggesting production of these precursors of monoterpenoids (geranyl-PP), diterpenoids and carotenoids (geranylgeranyl-PP) and sesquiterpenoids (farnesyl-PP). Terpenoids are commonly used by plants as defense against herbivores and many of them have antimicrobial properties (Mithöfer & Boland 2012). Furthermore, it has been shown that many bacteria are able to synthetize them (Yamada et al. 2014). However, to our knowledge there is no report on insect gut symbionts producing such compounds. A more plausible hypothesis is that geranylgeranyl-PP or farnesyl-PP might be taken up by the host and used as precursors for hormone synthesis. Geranylgeranyl-PP might be used to produce carotenoids as well. Carotenoids play a role in multiple physiological functions of the insect host (Heath et al. 2012; Sloan & Moran 2012). Host KEGG pathways having more DE enzymes in the hindgut wall, as compared to pocket were glycolysis/gluconeogenesis, pyruvate metabolism and pantothenate and CoA biosynthesis. Expression of these pathways, related to energy, CoA and acetyl-CoA production, is likely linked to the absorption of acetate and other short chain fatty acids through the hindgut epithelium (Bayon & Mathelin 1980; Terra & Ferreira 2009). Fatty acid biosynthesis KEGG pathways were DE as well, possibly related to high acetyl-CoA production rate in hindgut epithelial cells. Also KEGG pathways leading to synthesis of amino acids were DE in hindgut wall libraries (alanine, aspartate and glutamate metabolism, arginine biosynthesis and cysteine and methionine metabolism). This might be consequence of high availability of free ammonia and/or glutamate due to the degradation of insect waste nitrogen by hindgut wall bacteria (Alonso-Pernas et al. 2017). The KEGG pathway biosynthesis of antibiotics also showed higher number of DE mapped enzymes in hindgut wall, reflecting a possible expression of antimicrobial compounds in order to keep the symbiotic community under control (Garcia et al. 2010). However, the number of annotated contigs with assigned eukaryotic taxonomy is surprisingly low in hindgut wall compared to pockets (Table S1). This might be consequence of either low host RNA synthesis or masking of host’s transcripts by the highly abundant bacterial RNAs. In the latter case, it is possible that some host functions have been overlooked.

### 4.5. Differentially expressed (DE) metabolic pathways in pockets

In pocket libraries only a few KEGG pathways have more mapped enzymes of symbiotic origin compared to hindgut wall (Fig. 2). This might be a consequence of a lack of EC codes for many annotated enzymes, or might reflect that in second-instar larvae the pocket community is still ongoing a colonization phase and does not express yet all the particular functional pathways of the tissue. Two of the KEGG pathways DE by pocket symbionts were the TCA cycle and the synthetic pathway of protoheme groups from glutamate and Fe^2+^ (within porphyrin and chlorophyll metabolism). The expression of these pathways, together with HDE bacterial lipoyl synthase, supports occurrence aerobic metabolism and oxidative stress in pocket symbionts (Navari-Izzo et al. 2002; Girvan & Munro 2013). As mentioned before, no cellulases were DE in pocket, but enzymes processing oligo- and monosaccharides such as sucrose, fructose or maltose (within starch and sucrose metabolism) and glucose (within glycolysis/gluconeogenesis) were DE by pocket symbionts, suggesting that they might use the saccharides resulting from degradation of diet polysaccharides, such as cellulose and hemicellulose, in the hindgut. Pocket symbionts also DE more enzymes related to sulfur metabolism compared to hindgut wall. This, together with HDE insect sulfate transmembrane transporters in the pocket, suggest a role related to sulfur provisioning. Sulfur is needed for the synthesis of essential amino acids such as methionine and cysteine, the deficiency of which renders the insect vulnerable to plant defensive protease inhibitors (Broadway & Duffey 1986). In pockets, it appears that the insect differentially expresses all enzymes leading to eumelanin synthesis (within tyrosine metabolism pathway). This is in line with the host defensive deployment in this tissue already suggested by the analysis of HDE contigs (Nakhleh et al. 2017). An unexpected feature of the pockets is that the insect differentially expresses three enzymes of the KEGG pathway steroid hormone biosynthesis, one of which, the cholesterol monooxygenase (EC 1.14.15.6), is exclusive of the pathway. Steroid hormones may regulate the pocket cuticle synthesis as deduced from the HDE of insect cuticular proteins (Karlson 1989). Additionally, steroids might downregulate the synthesis of antimicrobial peptides, thus allowing symbiotic colonization (Gordya et al. 2016). Steroids are also involved in molting and metamorphosis (Niwa & Niwa 2016), supporting the abovementioned hypothesis of the pockets playing a role in host development. Insect chitin degrading and synthetizing enzymes were DE in pocket (within amino sugar and nucleotide sugar metabolism), in line with the pocket cuticular renovation or thickening previously discussed (Willis et al. 2012; Chapman 2013).

### 4.6. Conclusion

The present comparison of the gene expression of the pockets with the surrounding hindgut wall provided the first functional insight of these enigmatic structures. Combining the data of the present study with previous observations, we conclude that differences between the two tissues include, but may be not limited to: a) The active bacterial population in hindgut wall is mainly composed by anaerobes of the Firmicutes phylum, while those of the pockets is composed of aerobic and facultative anaerobes among which *Achromobacter* sp. is the most significant genus. b) The environmental origin of pocket symbionts is strongly supported by their taxonomy, their aerobic metabolism, their expression of transcripts related pathogen-like tissue colonization and the triggering of insect’s innate immunity. c) The environmental conditions between the two tissues appear to be remarkably different, the pockets having a higher oxygen concentration and oxidative stress, and lower bacterial competition and concentration of dietary toxins than the hindgut wall. d) The hindgut wall is likely to be the site of bacterial degradation of dietary recalcitrant polysaccharides and host nitrogenous wastes, while the pockets might play a role in stimulating host immunity, regulating host development and/or micronutrient absorption. e) The pocket bacterial community probably varies across larval stages as does that of the hindgut wall; therefore, its gene expression may change depending on larval maturity. Time-course RNAseq experiments may be useful to clarify this question.

## 5. Acknowledgements

The authors thank Dr. Heiko Vogel for his support during sequence analysis.

## Supplementary Material

The supplementary material of this manuscript can be downloaded at: https://www.dropbox.com/s/mmqmcoedvidyaic/Supplementary%20materials%20RNAseq.zip?dl=0

